# Gene gain and loss drive the diversification of *gig* immune genes in teleosts: structural and regulatory insights from Atlantic salmon

**DOI:** 10.1101/2025.07.01.662619

**Authors:** D. Manousi, S. Naseer, S.A.M. Martin, S.R. Sandve, M. Saitou

## Abstract

Interferon-stimulated genes (ISGs) are key players in vertebrate antiviral immunity. Among teleost ISGs, the grass carp reovirus-induced gene (*gig*) families 1 and 2 (*gig1* and *gig2*, respectively) are absent in mammals but conserved in fishes and amphibians, and they have been implicated in resistance to viral infections across several aquaculture species. In particular, *gig1* and *gig2* genes are transcriptionally induced by viral stimuli in teleosts such as zebrafish, grass carp, and salmonids, and recent studies have highlighted their potential involvement in resistance to economically important diseases like pancreas disease in Atlantic salmon. Yet, the rapid evolution of these genes hinders a comprehensive understanding of their diversification process and regulatory mechanisms. This study investigated *gig* gene evolution across teleosts, with a focus on Atlantic salmon (*Salmo salar*). Phylogenetic analysis across representative ray-finned fishes (Actinopterygii), including both teleosts and non-teleost outgroups such as the spotted gar (Holostei), indicated that gig1 is restricted to teleosts, with no identifiable homologs in non-teleost lineages. In contrast, gig2 genes are present in both teleosts and the spotted gar, suggesting an origin prior to the teleost-specific whole genome duplication (Ts3R), likely in early non-amniote vertebrates. Whole-genome duplication drove lineage-specific expansions, particularly of *gig2* in salmonids. Structural and transcriptomic analyses showed that *gig1* and *gig2* differ in domain composition, repeat content, and regulation. Our findings suggest the complex interplay of duplication history, structural divergence, and transcriptional regulation in shaping immune gene repertoires in teleosts, with implications for understanding host-pathogen interactions and aquaculture disease responses.

**Article summary:** This study investigates the evolutionary history and diversification of the gig immune gene families in aquatic species, with particular focus on Atlantic salmon. Phylogenetic and structural analyses revealed that gig1 and gig2 follow distinct evolutionary trajectories, shaped by whole-genome and tandem duplications. Further analysis of Atlantic salmon gig genes showed divergent structures and regulation, highlighting a general role of gig genes in antiviral Interferon-mediated immunity and additionally suggesting functional specialization across gig paralogs. Together, these findings improve our understanding of immune gene evolution in fishes and provide insights relevant to antiviral defense and disease management in aquaculture species.

## Introduction

Vertebrates have acquired complex immune systems to defend against a variety of pathogens (Iwama and Moran 2023). The interferon (IFN) signaling pathway plays a key role in vertebrate antiviral response as a molecular frontline in host defense (Dalskov *et al*. 2023). Upon viral infection, IFN activates a cascade of interferon-stimulated genes (ISGs), genes encoding proteins that suppress viral replication and enhance immune signaling and inflammatory responses (Walker *et al*. 2021). While insights into the functional diversification of ISGs have expanded, much remains to be explored outside the mammalian lineage (Purushotham *et al*. 2025).

Teleost fish, in particular, present an interesting group to study immune gene diversity and evolution, owing to their aquatic habitats, exposure to diverse pathogens, and history of multiple whole-genome duplications (Dornburg *et al*. 2021). Teleosts occupy environments ranging from freshwater rivers to deep-sea habitats, representing the most taxonomically and ecologically diverse group of vertebrates (Robertsen 2006; Sun *et al*. 2009; Mayer and Pšenička 2024). Their aquatic lifestyle imposes unique immunological challenges as they are continuously exposed to high pathogen loads via permeable surfaces such as gills, skin, and gut epithelium (Gomez *et al*. 2013; Wang *et al*. 2025). Teleosts possess a distinct immune system architecture, including specialized mucosal lymphoid tissues and the use of the pronephros for hematopoiesis, that differs markedly from that of terrestrial vertebrates. (Salinas *et al*. 2011; Levraud *et al*. 2019; Stosik *et al*. 2023). These ecological and physiological features are thought to have driven the diversification of ISG repertoires in teleosts. Notably, teleosts underwent an additional, teleost-specific whole-genome duplication (3R) (Amores *et al*. 1998; Taylor *et al*. 2003; Jaillon *et al*. 2004), with further duplications occurring in lineages such as salmonids and cyprinids (4R) (Macqueen and Johnston 2014; Blasweiler *et al*. 2023). These duplications provide opportunities for functional diversification of immune gene families, by potentially contributing to the expansion and functional specialization of immune genes (Glasauer and Neuhauss 2014; Clark *et al*. 2023) and allowing for adaptation to diverse pathogen pressures (Rivera and Swanson 2022; Tasnim *et al*. 2024).

Among the many interferon-stimulated gene (ISG) families in vertebrates, the grass carp reovirus-induced gene (*gig*) family stands out for being specific to fish and lower vertebrates, (including lampreys and amphibians), representing a lineage-restricted ISG family potentially shaped by aquatic pathogen pressures (Zhang 2003; Zhang and Gui 2004; Zhang *et al*. 2013; An *et al*. 2025). *Gig* genes show clear transcriptional responses to viral infection, across multiple species, including zebrafish, grass carp, and salmonids (Krasnov *et al*. 2011; Sun *et al*. 2014; An *et al*. 2025; Manousi *et al*. 2025). Consistently, in grass carp, overexpression of *gig1* and *gig2* enhanced resistance to grass carp reovirus (GCRV) (Sun *et al*. 2013). While some differences in the molecular responses of *gig* genes have been previously reported (Rao and Su 2015), their functional mechanisms and evolutionary relationships, especially between *gig1* and *gig2* genes, remain largely unknown. In addition, *gig* genes are expressed in response to economically important diseases in aquaculture species, such as cardiomyopathy syndrome (CMS) and pancreas disease (PD) in salmonids (Timmerhaus *et al*. 2012; Robinson *et al*. 2020; Krasnov *et al*. 2021). As a recent example, three tandemly duplicated *gig1* genes have emerged as candidate genes in a genome-wide association study for pancreas disease resistance in Atlantic salmon (Manousi *et al*. 2025). Collectively, these features suggest that *gig* genes are of particular interest due to their potential roles in aquaculture and disease resistance in aquatic vertebrates (Krasnov *et al*. 2011). While gig genes have received growing attention in the context of disease response and selective breeding, their evolutionary diversification and functional mechanisms are still incompletely understood. Though several studies have independently explored the evolutionary history of *gig1* and *gig2* (Zhang *et al*. 2013; Clark *et al*. 2023; An *et al*. 2025), only few have systematically focused on the functional diversification of *gig* gene families in response to their abundance. To address this gap, a more integrated approach is needed to understand how gene duplication, expression patterns, and functional divergence have shaped the evolution of *gig* genes.

In this study, we present a comprehensive investigation of *gig* gene evolution and functional diversification in teleosts, with particular focus on Atlantic salmon. Atlantic salmon (*Salmo salar*) provides an interesting system to investigate gig evolution and regulation; occurrence of a lineage-specific (Ss4R) whole-genome duplication in the salmonid ancestor led to expansion of the gene repertoire, facilitating functional divergence and specialization. At the same time, Atlantic salmon is one of the fish species with the most developed genomic resources, including large transcriptomic and chromatin accessibility datasets generated in the AquaFAANG project (https://www.aqua-faang.eu/) (Johnston *et al*. 2024). We integrate phylogenetic analysis and functional genomics data from in-vivo and in-vitro viral challenges to elucidate how *gig1* and *gig2* have evolved and contributed to immune gene innovation in non-mammalian vertebrate lineages.

## Materials and Methods

### Species selection and gene family phylogenetic tree construction

To investigate the evolutionary history and diversification of *gig* genes across teleosts, we selected 11 representative species and constructed a gene family phylogenetic tree. All genome assemblies and gene annotations were retrieved from the Ensembl database (version 112), ensuring consistency in gene models and facilitating reliable cross-species comparisons of *gig* homologs. Species were chosen to represent major teleost clades and to include lineages that have undergone whole-genome duplication (WGD) events, which are relevant to the expansion and diversification of immune-related gene families. The selected taxa, Atlantic salmon (*Salmo salar*), Northern pike (*Esox lucius*), zebrafish (*Danio rerio*), common carp (*Cyprinus carpio*), Coho salmon (*Oncorhynchus kisutch*), Chinook salmon (*Oncorhynchus tshawytscha*), rainbow trout (*Oncorhynchus mykiss*), brown trout (*Salmo trutta*), Japanese medaka (*Oryzias latipes*), three-spined stickleback (*Gasterosteus aculeatus*), and spotted gar (*Lepisosteus oculatus*) (**Table 1**), span a range of phylogenetic positions (e.g., Otophysi, Euteleostei, Holostei) and encompass both WGD and non-WGD lineages.

**Table 1.** Genome assemblies utilized in this study.

| Scientific name | Common name (Ensembl) | Accession | N50 | Citation |
| --- | --- | --- | --- | --- |
| *Oncorhynchus tshawytscha* | Chinook salmon | GCA_01829614 5.1 | 2884253 | (Christensen *et al.* 2018) |
| *Oncorhynchus mykiss* | Rainbow trout | GCA_01326573 5.3 | 39165350 | (Gao *et al.* 2021) |
| *Gasterosteus aculeatus* | Stickleback | GCA_01692084 5.1 | 20445003 | (Nath *et al.* 2021) |
| *Lepisosteus oculatus* | Spotted gar | GCA_00024269 5.1 | 6928108 | (Braasch *et al.* 2016) |
| *Cyprinus carpio* | Common carp<br>german mirror | GCA_90522157 5.1 | 28473900 | (Blasweiler *et al.* 2023) |
| *Danio rerio* | Zebrafish | GCA_00000203 5.4 | 4737936 | Genome reference consortium |
| *Oncorhynchus kisutch* | Coho salmon | GCA_00202173 5.2 | 68265700 | - |
| *Salmo salar* | Atlantic salmon | GCA_90523706 5.2 | 28058890 | - |
| *Salmo trutta* | Brown trout | GCA_90100116 5.1 | 52209666 | (Hansen *et al.* 2021) |
| *Esox lucius* | Northern pike | GCA_01100484 5.1 | 37473588 | - |
| *Oryzias latipes* | Japanese medaka HNI | GCA_00223467 5.1 | 31218526 | (Ichikawa *et al.* 2017) |

To identify *gig* homologs, the coding DNA sequences of the originally reported *gig1* and *gig2* genes were retrieved (Zhang and Gui 2004) and used as queries against the selected genomes. The corresponding gene names and descriptions were retrieved via BioMart (Ensembl v112) and NCBI gene annotations when applicable (**Supplementary Table S1**). For each gene, the longest protein-coding transcript was retained, and its amino acid sequence was used for downstream analysis.

Amino acid sequences were aligned using GUIDANCE2 in order to assess the reliability of sequence alignments and filter out poorly aligned regions (Sela *et al*. 2015). Alignments were refined by excluding positions with consistency scores below 0.93 (site alignments present in > 93% of GUIDANCE2 replicates), ensuring robust phylogenetic inference. Reliability filtering of sequence alignments removed one Atlantic salmon *gig2* ortholog, ENSSSAG00000082930, which could be a pseudogene or partial duplication. The remainder alignments were used to construct maximum likelihood phylogenies in IQ-TREE version 2.3.3, implementing the *Model-finder* option in order to detect and utilize the best-fit substitution model for each *gig* phylogeny (Nguyen *et al*. 2015; Kalyaanamoorthy *et al*. 2017). Support values for the two *gig* trees were estimated using the ultrafast bootstrap method implemented in IQ-TREE and 1,000 replicates (Hoang *et al*. 2018). The resulting trees, together with a species tree constructed in timetree.org, were visualized in the iTOL platform (Letunic and Bork 2021). Phylogenetic clades in the constructed gene trees were then defined based on the visualized phylogenies.

### *Gig* functional domain prediction

To characterize conserved protein domains and infer functional divergence between *gig1* and *gig2* genes, the predicted amino acid sequence of the longest isoform per gene was analyzed using InterProScan v98.0 (Paysan-Lafosse *et al*. 2023). This tool integrates domain predictions from multiple databases, including Pfam, SMART, and Gene3D. Domains found in fewer than 5% of all *gig* genes within a given clade, typically appearing only 2–3 times, were excluded to reduce the influence of rare features. By doing so, we aimed to improve signal specificity in detecting conserved functional domains across teleost lineages. In addition, to decrease the dimensionality of the dataset, filtered domains identified across multiple databases were clustered together based on functional similarity, allowing assessment of potential divergence in molecular function between *gig1* and *gig2* genes. Domain annotation and clustering were performed based on established definitions from the literature, in particular for predicted domains related to ADP-ribosylation (Di Girolamo and Fabrizio 2018). Alternatively, in cases such as signal peptide and membrane bound protein-related regions, subdomains were grouped under a single functional category (e.g., ‘signal peptide region’ and ‘membrane-bound protein region’, respectively), reflecting shared biological roles (**Supplementary Table S2**). To visualize functional domains in Atlantic salmon *gig* genes, sequentially located features that belonged in the same functional cluster were collapsed together in a single feature.

### Detection of repeat content within coding regions of *gig* genes

To identify internal sequence repetitions within *gig* paralogs, the nucleotide sequences were analyzed using a self-comparison approach. Pairwise comparisons of the coding sequence for each *gig* gene were performed using the ‘dotPlot’ function in the R package *seqinr* to visualize self-aligned sequence and detect repetitive motifs (Charif and Lobry 2007; R Core Team 2021). A sliding window approach was implemented to detect local similarities within each sequence, using a window size of 10 nucleotides and a step size of a single nucleotide. A dot was plotted at each position where 9 or more nucleotides matched between two overlapping windows. This allowed for the visualization of internal repeats and low-complexity regions that may indicate recent duplications or structural features relevant to *gig* gene evolution.

### Evolutionary characterization of Atlantic salmon *gig* genes

Rediploidization timing has a large impact on sequence similarity, with earlier occurrences allowing for higher sequence divergence. To investigate the genomic context and evolutionary origins of *gig* genes in Atlantic salmon, we examined the genes’ physical positions in relation to known patterns of rediploidization following the Ss4R duplication event.

To achieve this, chromosomal regions reflecting early (Ancestral Ohnolog Resolution, AORe), and delayed (Lineage-specific Ohnolog Resolution, LORe) phases of post-duplication divergence (Lien *et al*. 2016; Gundappa *et al*. 2022) were retrieved from the Salmobase browser (https://salmobase.org/). The retrieved information were then compared against the physical coordinates of gig genes to investigate potential links between *gig* paralogs and rediploidization timing (**Supplementary Table S3**).

In addition to genome-wide duplication, we also assessed the potential contribution of tandem duplication events to *gig* gene family expansion. Tandemly duplicated genes were defined as those located on the same chromosome and separated by less than 1 Mbp. This classification allowed us to distinguish between large-scale (WGD-derived) and local duplication mechanisms in shaping the current genomic architecture of the *gig* family in Atlantic salmon.

### Manual annotation of *gig* paralogs

Many *gig* genes across species showed substantial sequence divergence. This was reflected in the large number of amino acid residues removed during alignment filtering. We hypothesized that this pattern reflects genuine biological variation, especially a signature of rapid evolution in immune-related genes, rather than a result of annotation errors. To test this hypothesis, we examined the transcript structures of *gig* genes using proprietary and publicly available RNA-seq datasets. While this phenomenon was not limited to Atlantic salmon, we focused on this species because of the rich availability of both short– and long-read transcriptomic data.

Short-read RNA-seq data from poly(I:C) stimulation and SAV3 infection challenges were obtained from the AQUAFAANG public repository (accession number PRJEB50076, Clark et al., 2023) and the European Nucleotide Archive (accession number PRJEB85594, Hillestad *et al*. 2020), respectively. RNA-seq reads were aligned to the Atlantic salmon genome (Ssal_v3.1, Ensembl v112) using STAR 2.7.10a (Dobin *et al*. 2013) and the alignment coverage of samples with the highest *gig* gene expression was visualized using the Integrative Genome Viewer (IGV) (Robinson and Zemo jtel 2017).

In addition to short-read RNA-seq data, long-read transcriptomic data from 12 healthy individuals, previously sequenced using Oxford Nanopore Technologies (ONT) were additionally incorporated to refine gene annotations and assess the limitations of short-read mapping in complex genomic regions. Details regarding sampling and alignment procedures for the ONT-sequenced individuals against the Ssal_v3.1 genome are available elsewhere (Sandholm 2022). Aligned transcriptomic data from both platforms were compared to the existing gene annotation (Ssal_v3.1) in IGV, enabling refinement and validation of exon-intron boundaries, identification of transcription start sites (TSS), and confirmation of alternative isoforms (**Supplementary Table S4**).

To further evaluate coding potential, sequences corresponding to candidate exons (including 100 bp upstream and downstream flanking regions) were extracted and analyzed using the NCBI ORFfinder. Where necessary, predicted protein products were submitted to InterProScan (Paysan-Lafosse *et al*. 2023) to confirm the presence or absence of conserved functional domains.

### Discovery of interferon-specific regulatory regions using chromatin accessibility and histone modification data

Interferon-stimulated response elements (ISREs, (Collet and Secombes 2001) were examined within 2 kb upstream and 200 bp downstream of the transcription start site of all *gig* genes to assess their regulatory potential in interferon-mediated immune responses (Holen *et al*. 2023). To evaluate the functional relevance of these predicted regulatory elements, publicly available chromatin accessibility (ATAC-seq) and histone modification (H3K27ac, H3K27me) data were incorporated. These datasets were derived from an interferon-stimulation experiment in Atlantic salmon head kidney tissue, where poly(I:C)-treated samples were compared to unstimulated controls (Clark *et al*. 2023). In brief, DNA was extracted from head kidney tissue of six individuals per condition, and sequencing was performed using ATAC-seq and ChIP-seq technologies. To assess signatures of chromatin accessibility and histone modifications, the nf-core pipelines *atacseq* and *chipseq* were used, providing standardized modular workflows for quality control, read alignment, and peak calling (Ewels et al. 2020).

Further refining the regulatory element identification, an Irreproducible Discovery Rate (IDR) analysis was performed using the software ChIP-R (Newell *et al*. 2020) to ensure high reliability of regulatory signals across samples in each condition and reduce false positives. Finally, the differential accessibility of peaks between conditions was assessed using DESeq2 (Love *et al*. 2014) and regulatory elements were considered statistically significant if they exhibited an FDR-adjusted p-value < 0.05. These analyses enabled the identification of ISRE motifs and chromatin modifications that potentially regulate *gig* gene expression in response to interferon stimulation.

### Transcriptomics data and gene expression analysis of Atlantic salmon *gig* genes

To explore the immune-related functional divergence of *gig1* and *gig2* in Atlantic salmon, we analyzed transcriptomic responses to both interferon stimulation and viral infection using three publicly available RNA-seq datasets. Specifically, two datasets were derived from interferon-stimulation experiments (ENA accession number PRJEB50076, Clark *et al*. 2023). In these studies, Atlantic salmon head kidney tissue and primary head kidney cells were treated with poly(I:C), while control groups received PBS medium. The third dataset (ENA accession number PRJEB85594) originated from a viral challenge experiment involving Salmonid alphavirus (SAV3), in which heart tissue was sampled from fish that survived or died following 11 days of infection (Hillestad *et al*. 2020). For all datasets, total RNA was extracted and sequenced using Illumina-based RNA-seq platforms. Quality filtering, alignment of sequencing reads to the salmonid transcriptome (Ssal_v3.1, GCA_905237065.2), and quantification of gene expression were performed using the nf-core *rnaseq* pipeline (Patel *et al*. 2024).

Differential gene expression analysis was performed independently for each of the three datasets using DESeq2 (Love *et al*. 2014), with a threshold of FDR-adjusted p-value < 0.05.

### Use of a Large Language Model

We have used a Large Language Model (LLM) in our work, specifically ChatGPT (GPT-4) developed by OpenAI. The model was accessed through the OpenAI API, and our interactions primarily involved two types of queries: seeking assistance with coding-related issues and refining English expressions as non-native speakers. Our queries were structured in natural language, with iterative refinements applied when necessary to improve clarity and accuracy. As authors, we acknowledge our responsibility for the accuracy of the content generated, ensuring that it is free from plagiarism, and properly attributes all sources, including material produced by the LLM.

## Results

### Expansion and divergence of *gig1* and *gig2* gene families across teleosts species

Comparative genomic analysis across ancestral and teleost species revealed distinct evolutionary trajectories for *gig1* and *gig2* gene families (Figure 1). Consistent with previous reports (Zhang et al. 2013), *gig2* genes were identified in all surveyed non-amniote vertebrates. In contrast, *gig1* genes were exclusively identified in teleosts. Notably, we did not identify any *gig1* homologs in the spotted gar genome using our criteria. However, a recent study (An *et al*. 2025) reported a candidate *gig1* gene in spotted gar based on conserved domain architecture. Upon our inspection, the gene in question (ENSLOCG00000006921) did show similarities in domain composition, but in Ensembl, it clusters separately in its own orthogroup (ENSGT00940000174456). Given this ambiguity, and to ensure consistency in phylogenetic comparison, we excluded this gene from the *gig1* set used in our study. Lastly, comparing gene family sizes between species that experienced lineage-specific WGD events (e.g., common carp, salmonids) and their closest non-duplicated relatives (e.g., zebrafish, northern pike) suggests that whole-genome duplications are associated with expansion of the two *gig* families, but with clear lineage-specific differences; *gig1* genes underwent expansion following the common carp WGD (Xu et al. 2019), whereas expansion of the *gig2* genes seems to have occurred after the salmonid WGD (**Figure 1**).

**Figure 1.**
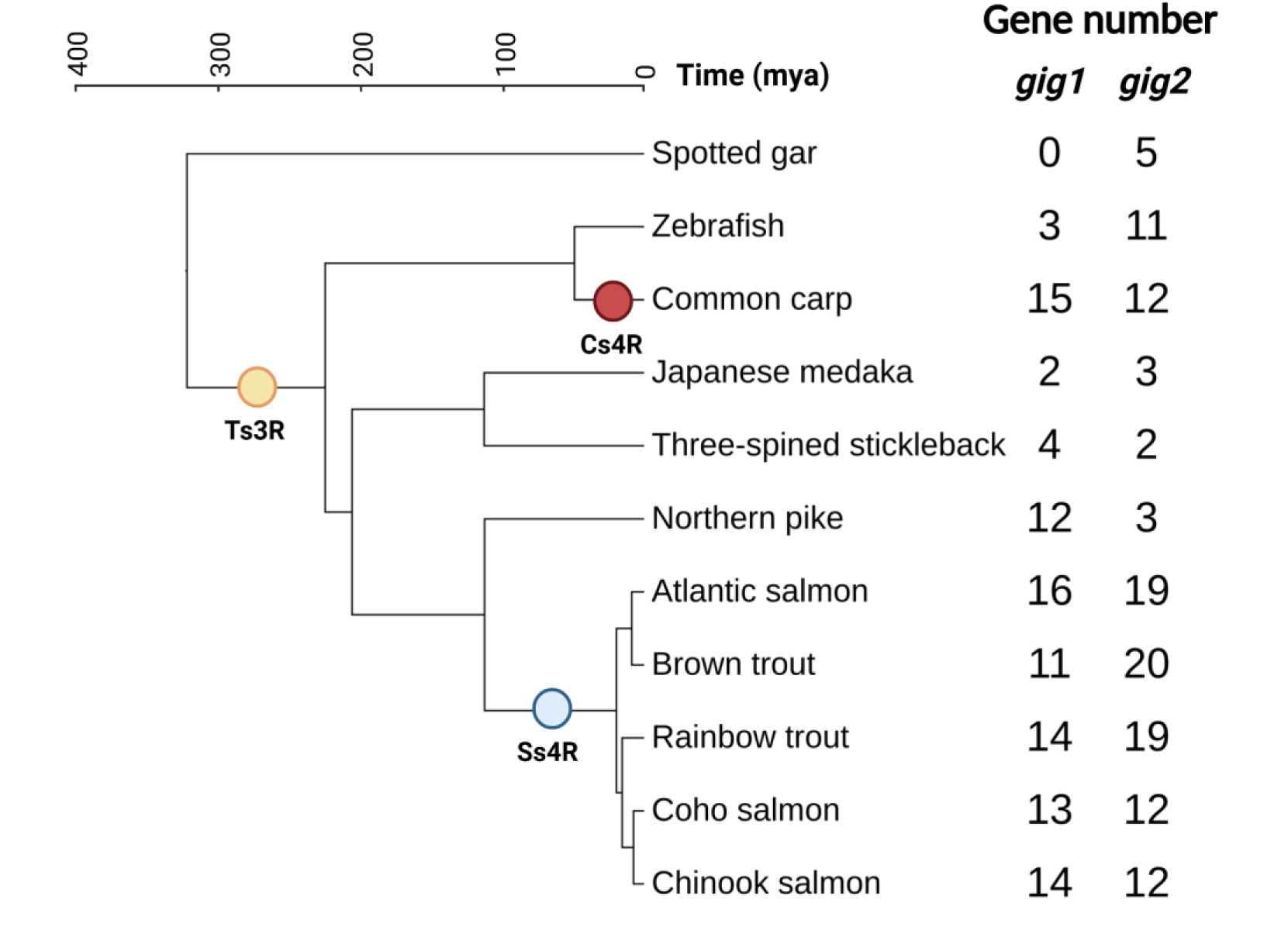
Gig1 and gig2 gene abundance across teleost species and the outgroup species spotted gar (Lepisosteus oculatus). A phylogenetic tree for 11 species was constructed in Timetree.org. Yellow, red and blue annotations represent the teleost-specific (Ts3R), carp-specific (Cs4R) and salmonid-specific (Ss4R) whole genome duplication events, respectively. Tree scale shows the chronological divergence of species in million years (mya). Created in BioRender. Manousi, D. (2025) https://BioRender.com/rz4gexa

To further investigate the evolutionary history of *gig* genes, including the temporal patterns and role of different duplication mechanisms in gene family expansion events, we constructed phylogenetic trees using the protein translation of the longest coding transcripts (**Figure 2 and Supplementary Figures 1 and 2**). Multiple sequence alignments were refined using GUIDANCE2 (Sela et al. 2015) to eliminate potential annotation errors and excessively divergent residues (Materials and Methods). Sequence reliability filtering revealed that 31% of *gig1* alignments were retained after filtering low-conservation sites, compared to 49% for *gig2*. These results indicate a higher degree of sequence conservation in *gig2* compared to *gig1*. Additionally, one Atlantic salmon *gig2* gene (ENSSSAG00000082930) was removed during the filtering process due to extreme sequence divergence, possibly linked to pseudogenization. The phylogenetic analysis indicated two major clades of *gig1* genes (*gig1A* and *gig1B*) and three (*gig2A*, *gig2B*, and *gig2C*) clades in *gig2* genes (**Figures 2a** and **2b and Supplementary Figures 1 and 2**). Further investigation of the phylogenetic structure also supported that the *gig1B* clade underwent expansion in common carp, likely driven by both WGD and tandem duplications; the common carp genes clustered in *gig1B* clade are located on two scaffolds in the carp genome assembly (scaffolds CAJNDQ010000023.1 and CAJNDQ010000028.1).

**Figure 2.**
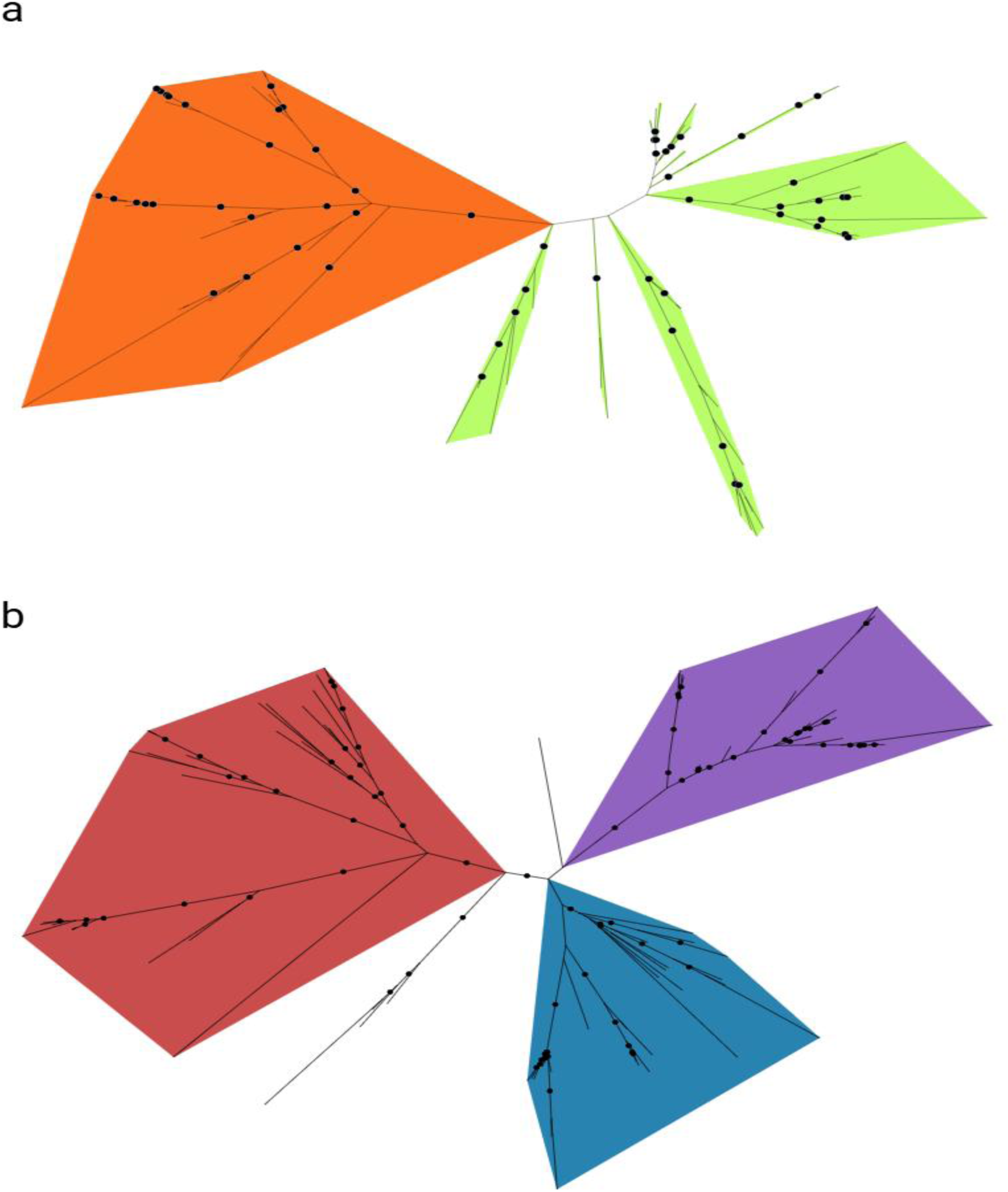
Unrooted phylogenetic trees of gig1 (top) and gig2 (bottom) genes across 11 species. Ortholog gig genes across species were retrieved from the Ensembl annotation gene gain/loss trees. The protein translation of the coding sequence for the longest transcript was aligned using GUIDANCE2 and unreliable residues were removed based on the default thresholds (0.93). Phylogenetic trees were built in IQ-TREE v2.3.3 using the alignments and 1000 replicates to calculate bootstrap values, shown as filled circles for values ≥ 80%. The ‘Model Finder’ option was used to detect the best-fit substitution model for each tree. 2a. Sequence divergence within the gig1 family suggests one possible split in the middle of the tree forming two separate clades, hereby named gig1A (orange) and gig1B (green). 2b. Sequence divergence within the gig2 family suggests three possible splits in the middle of the tree forming two separate clades, hereby named gig2A (red) and gig2B (blue) and gig2C (purple). Created in BioRender. Manousi, D. (2025) https://BioRender.com/iaw38fk

In contrast, *gig2* genes exhibited more complex, lineage-specific divergence. While *gig2A* and *gig2B* clades expanded in common carp, *gig2C* showed a pronounced increase in gene duplicates among salmonids. Within salmonids, *gig2* continued to diversify, with brown trout (*Salmo trutta*) showing expansion of ancestral-like *gig2A* genes, whereas rainbow trout exhibited an increase in *gig2C* **(Supplementary Figure 2)**. Notably, brown trout exhibited an expansion of ancestral-like *gig2A* genes, whereas rainbow trout showed an increase in *gig2C* copies. These lineage specific patterns of *gig2* expansion events may reflect neutral gene family size evolution or species-specific adaptations to environmental and/or pathogenic pressures, or a combination of these hypotheses. However, the functional implications of these evolutionary trajectories, including how regulatory and structural divergence has shaped the immunological roles of *gig* gene paralogs, remain to be clarified.

In the following sections, we focus on the Atlantic salmon and explore in detail genomic context, duplication history, and transcriptional regulation to uncover how gene duplication has contributed to the diversification of *gig* gene function across teleost species. For that reason we selected all gig genes detected in Atlantic salmon based on the Ensembl database and classified them based on phylogenetic clade and physical position on the salmonid genome (Ssal_v3.1, **Supplementary table S4**)

### Genome duplication and tandem repeats shaped the *gig* gene emergence and structural changes in Atlantic salmon

Atlantic salmon harbors the highest number of *gig* genes among the surveyed species (**Figure 1 and Supplementary Table S4**), a pattern likely shaped by the salmonid-specific (Ss4R) whole genome duplication that occurred approximately 125 million years ago (Gundappa *et al*. 2022). For clarity and reproducibility, we assigned simplified names to Atlantic salmon gig paralogs based on three criteria: their phylogenetic clade (e.g., *gig1A, gig2B*), chromosomal location (e.g., ssa02), and relative physical order (e.g., *a, b, c*). For instance, *gig2B_ssa02d* refers to the fourth identified *gig2B* gene located on chromosome ssa02. This naming is used throughout the manuscript and figures. A complete mapping between these names and Ensembl gene IDs is provided in **Supplementary Table S4**.

To investigate the evolutionary origin and structural divergence of *gig* genes in Atlantic salmon, we examined chromosomal organization, duplication patterns, and predicted protein features. We obtained synteny information from the latest version of the Atlantic salmon genome (ssal_v3.1) using the Salmobase browser (https://salmobase.org/) and assessed the physical location of *gig* paralogs considering the impact of the Ss4R genome duplication.

The salmonid genomes experienced delayed rediploidization in distinct regions of the genome. Genomic regions are therefore subdivided into AORe (early rediploidization; Ancestral Ohnolog Resolution) and LORe (late rediploidization; Lineage-specific Ohnolog Resolution) (Gundappa *et al*. 2022). Rediploidization history impacts sequence divergence as well as the probability of functional divergence; synteny analysis using data from Salmobase (Ssal_v3.1) revealed that three *gig1* and one *gig2* genes are located in AORe regions, while the remaining 13 *gig1* and 15 *gig2* genes reside in LORe regions (**Figure 3**). Two *gig2* genes were located on unplaced scaffolds (CAJNNT020001543.1 and CAJNNT020001561.1) and were therefore excluded from further analyses. Our results suggest that the majority of the 32 Atlantic salmon *gig* genes expanded following salmonid lineage diversification and distributed across 10 chromosomes. A high proportion of them was tandem duplicated, particularly within LORe regions on chromosomes ssa03, ssa04 and ssa08 (**Figure 3**), but no tandem duplicates were identified within AORe regions.

**Figure 3.**
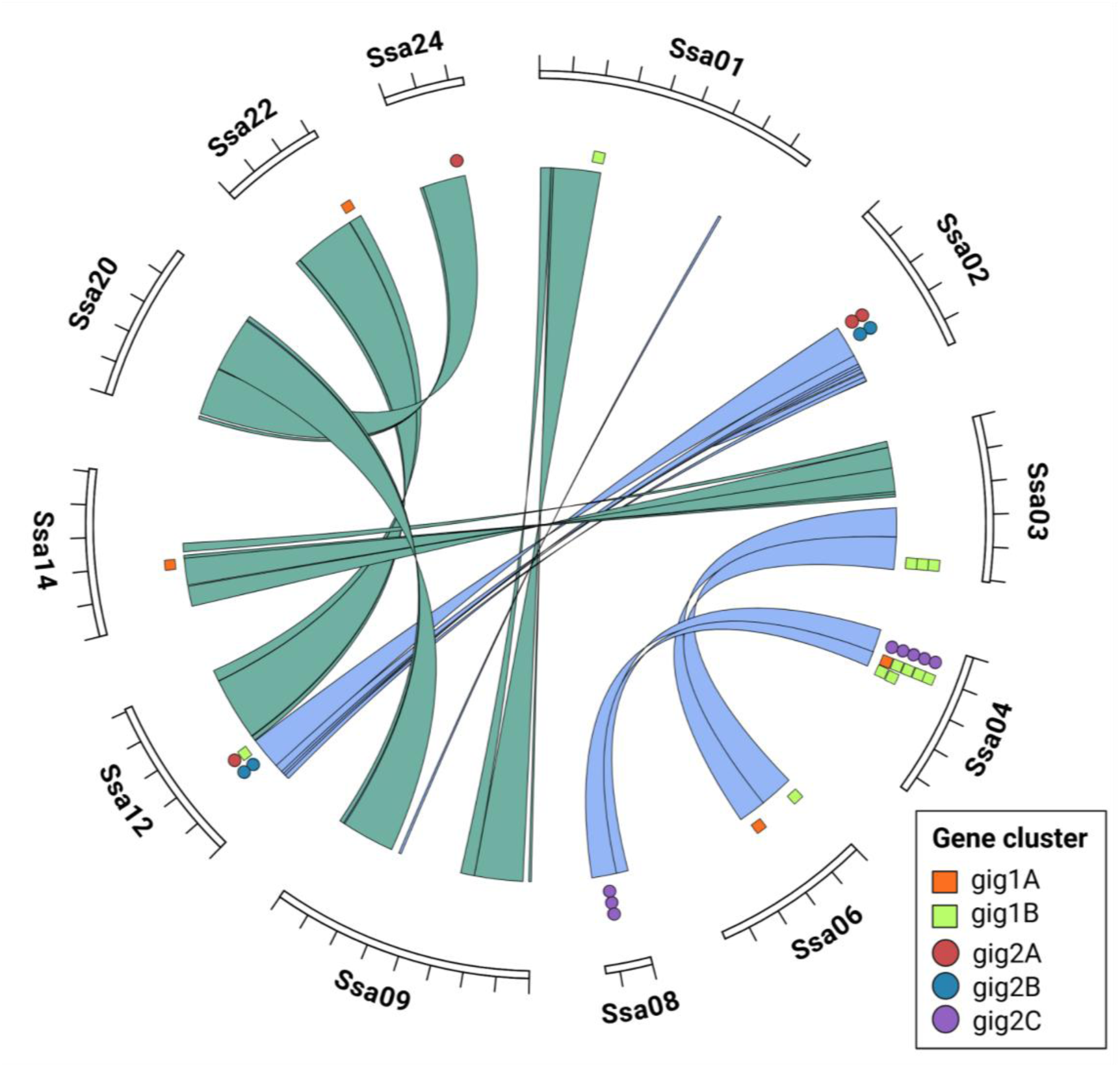
Circos plot of the physical location of gig1 (square) and gig2 (circle) genes in the salmonid genome. Colored shapes represent gene clusters based on phylogenetic analyses. Within the center of the plot, green ribbons link together genomic regions that underwent re-diploidization early after the Ss4R WGD event, (∼100mya, ancestral-specific ohnolog resolution, AORe) while blue ribbons connect the regions that underwent late re-diploidization within the last 50mya (lineage-specific ohnolog resolution – LORe) (Gundappa et al. 2022). Created in BioRender. Manousi, D. (2025) https://BioRender.com/8qiyb8b

Protein domain prediction using InterPro (version 102.0, Paysan-Lafosse *et al*. 2023) identified 17 conserved domains across all Atlantic salmon *gig* genes, which were grouped into eight clusters based on the literature research as well as functional similarity of predicted domains retrieved from different databases (**Figure 4 and Supplementary Table S2**). Notably, *gig1* genes contained exclusively ‘*signal peptide’-*related as well as ‘*SI:CH211-198C19.1-RELATED’* domains, with the latter domain being detected across all *gig1* genes, suggesting a high degree of structural conservation within this family. Similarly, *gig2* genes obtained exclusively the ‘*Gig2-(like)’* protein, ‘*ADP-ribosylation*’-related and ‘*coil’* domains, with the latter features additionally overlapping ‘*consensus disorder*’ predicted domains (**Figures 4a** and **4b**).

**Figure 4.**
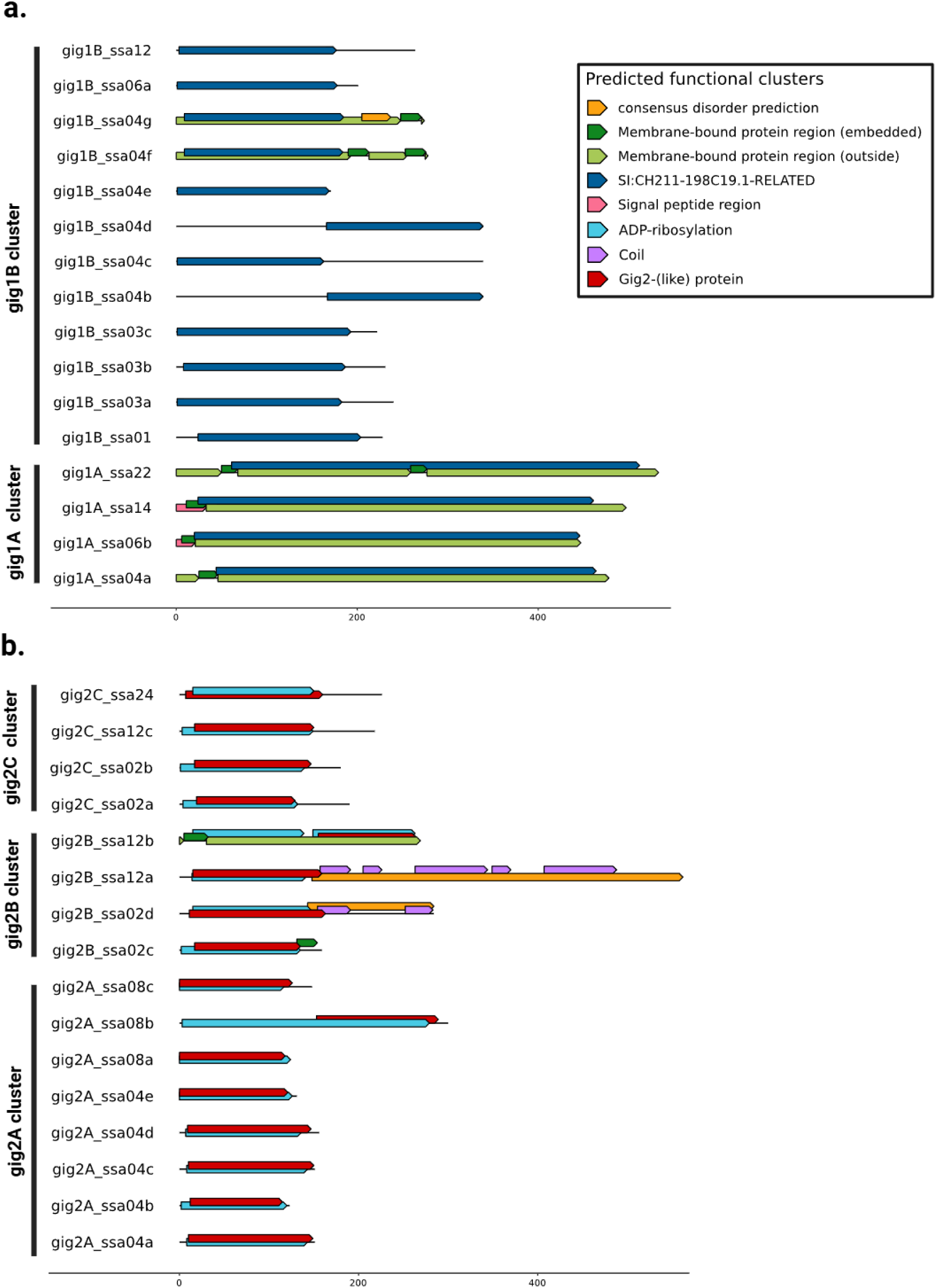
Predicted domains visualized for Atlantic salmon gig1 (a) and gig2 (b) ortholog genes. Colored bars and arrows indicate the functional group of a predicted domain whereas salmonid ortholog names are highlighted in respect to their phylogenetic clade. Black lines distinguish gene clusters as those were defined based on phylogenetic analysis. Created in BioRender. Manousi, D. (2025) https://BioRender.com/x8fy7hn

This gene family-specific domain composition supports the functional specification between *gig1* and *gig2*. Furthermore, there was variation in the length of the *‘SI:CH211-198C19.1-RELATED’* domain between *gig1A* genes and *gig1B* genes (**Figure 4a**), pointing to potential functional divergence within the *gig1* family. Besides *gig* family-specific domains, membrane-related and *consensus disorder* domains were found across genes of both families (**Supplementary Table S2**). Among *gig1* genes, six carried membrane-related domains and four carried ‘*signal peptide*’ domains; these were mostly located on ssa06, ssa04, ssa14, and ssa22. Meanwhile, only two *gig1B* genes, *gig1B_ssa04f* and *gig1B_ssa04g*, which are tandem duplicates, retained such domains (**Figure 4a**).

Functional diversification within gene families has been known to occur through repeat motif expansions (Pajic *et al*. 2022). We therefore examined the presence of repeat structures within *gig* genes. Comparison of the self-alignments for *gig1* and *gig2* genes (**Figure 5 and Supplementary Figures 3-6**) identified several different tandem repeats and fusions, in accordance with earlier reports (Zhang *et al*. 2013). Repeat content was detected in genes located within large tandem duplication clusters; *gig1* genes located on chromosome ssa04, and *gig2* genes located across chromosomes ssa02, ssa08, and ssa12. Interestingly, *gig1* genes with elongated domains lacked repeats, whereas *gig2* genes containing heavily repeated sequence exhibited elongated and duplicated ADP-ribosylation domains (**Figure 5, Supplementary Figures 1 and 2**). In addition, two WGD-derived duplicate genes with high repeat content, namely *gig2B_ssa02d* and *gig2B_ssa12a,* overlapped predicted *consensus disorder* and *coil* domains (**Figures 4b** and **5**). While the particular genes contained an exceptionally high abundance of glutamate (E) repeats in their amino acid sequences, there were not substantial differences in the length of their respective *Gig2 protein*-related domain. However, besides this extensive repeat expansion, the overall length of the core Gig2-related domain remained relatively stable across *gig2* paralogs. This suggests that while the central functional domain may be conserved, structural variation has occurred primarily in the N-terminal end of the protein. Although the functional consequences of these changes remain to be determined, they raise the possibility of differential regulatory or immunological roles among paralogs. Taken together, these findings indicate that *gig* gene expansion in Atlantic salmon has been shaped not only by large-scale duplication events but also by within-gene tandem repeat expansions and structural rearrangements, such as internal repeat proliferation. To assess whether such gene-specific structural features correspond to functional divergence, we next examined the transcriptional responses of *gig* paralogs to immune stimulation.

**Figure 5.**
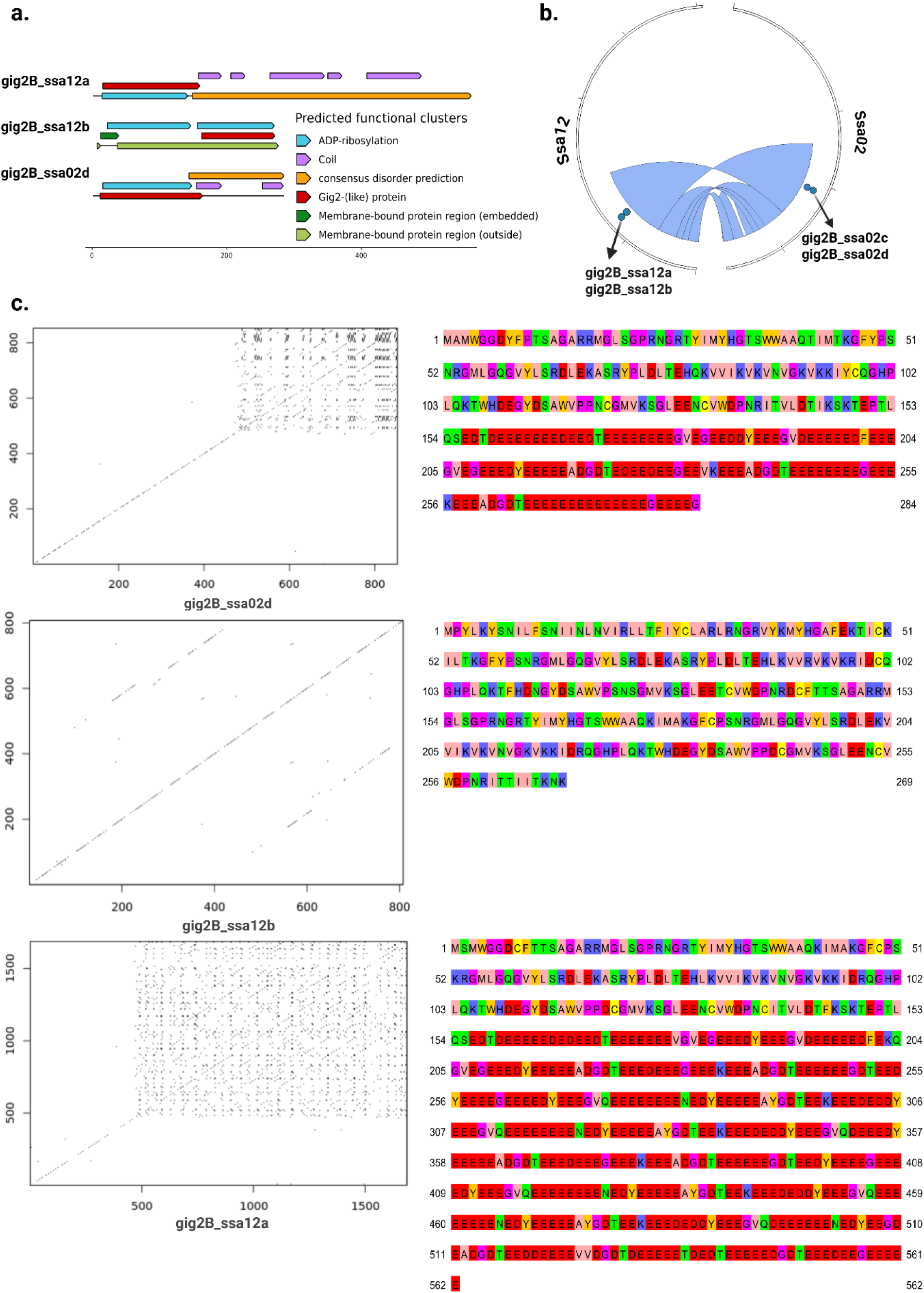
Repeat content and predicted functional domains for gig2 paralogs located on the gig2B phylogenetic clade. (a.) Functional domain prediction based on the amino acid translation of gig coding sequence. (b.) Physical position of paralogs in the salmonid genome. Blue ribbons indicate chromosomal regions of late rediploidization (LORe). (c.) On the left, the self-alignment of the coding sequence for gig2 paralogs is shown while on the right, the amino acid translation of the aligned sequence for the respective genes is displayed. Created in BioRender. Manousi, D. (2025) https://BioRender.com/4f0vqtr

### Expression divergence of *gig* paralogs reveals functional diversification under immune stimulation

To explore the immunological roles of *gig* genes in Atlantic salmon and distinguish between interferon-induced gene regulation and more complex host responses to viral infection, we analyzed their expression responses to antiviral stimulation using three independent RNA-seq datasets. These included *in vitro* and *in vivo* poly(I:C) stimulation of head kidney (HK) cells and tissues (Clark *et al*. 2023), and heart tissue transcriptomes from a salmonid alphavirus (SAV3) infection survival challenge (Hillestad *et al*. 2020).

*Gig1* genes displayed heterogeneous transcriptional responses, with expression patterns differing between the gig1A and gig1B clades (**Figure 6**). Several *gig1B* genes, particularly those located on chromosomes ssa03 and ssa06, consistently responded to both *in vitro* and *in vivo* immune stimulation, including SAV3 infection. In contrast, only one out of four *gig1A* genes, *gig1A_ssa14*, showed notable upregulation under poly(I:C) treatment. This gene also exhibited relatively high baseline expression, suggesting it may play a more prominent immune role within the gig1A clade. Other gig1A members showed minimal or no transcriptional induction under any condition tested. On ssa04, multiple gig1B paralogs were detected; however, only *gig1B_ssa04b* and *gig1B_ssa04g* responded to stimulation, with the latter showing a more pronounced response in head kidney tissue. Overall, the expression profiles of *gig1* genes did not correlate strongly with phylogenetic clade structure, indicating that post-duplication regulatory divergence has played a key role in shaping their transcriptional pattern.

**Figure 6.**
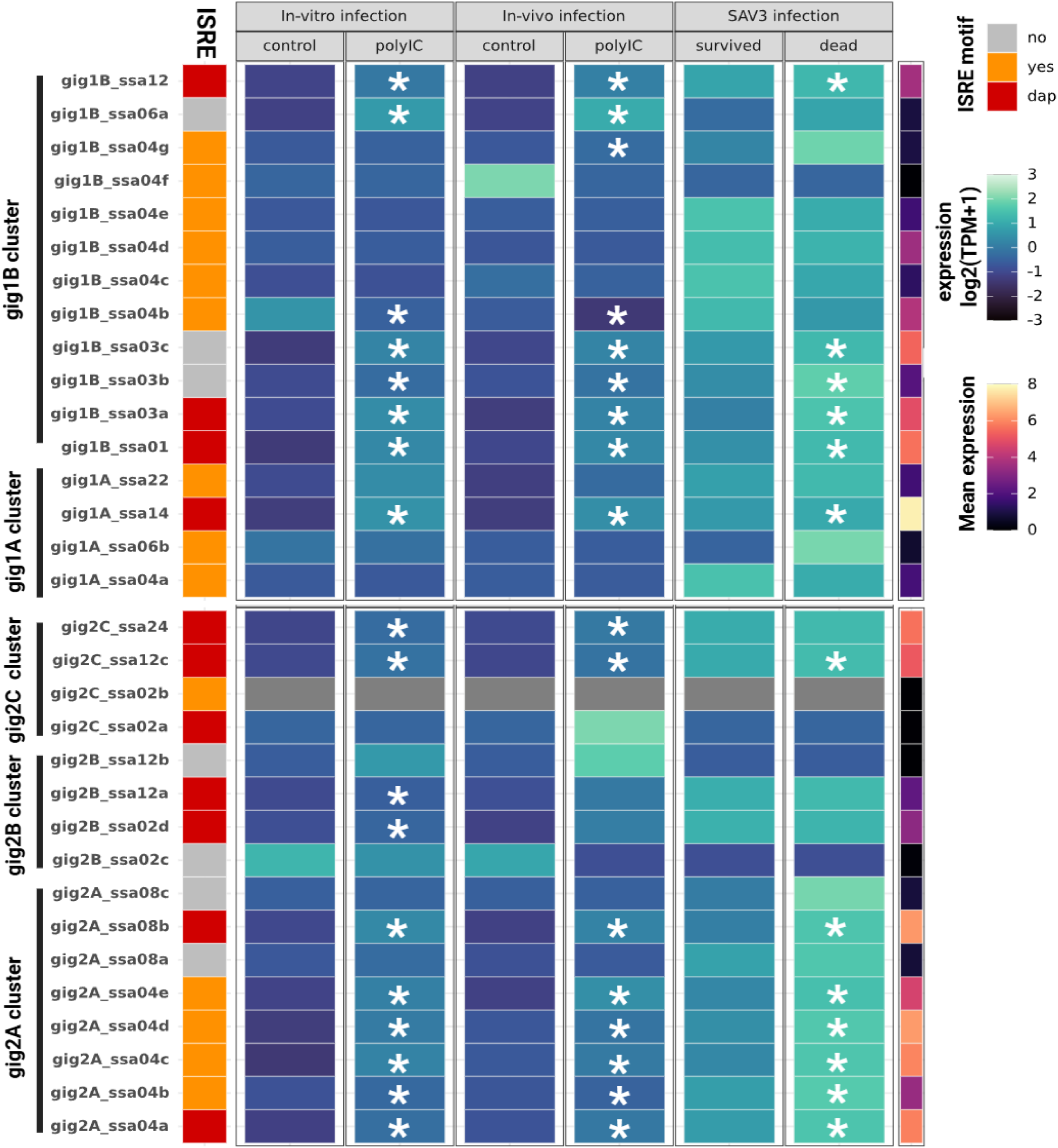
Comparative transcriptomics analysis of gig1 and gig2 genes in Atlantic salmon. Gig genes are sorted based on phylogenetic clustering. The expression panel shows the mean gene expression of gig genes across head kidney samples treated in vitro (cells) and in vivo (tissue) with poly(I:C) in addition to heart tissue transcriptome data from samples challenged with PD disease (SAV3 virus) in an 11-day survival trial. Colored boxes indicate mean gene expression of sample replicates in the range of 0 to 8 log2(TPM + 1) values. A discrete color annotation is used to indicate genes with at least one detected IFN stimulation response element (ISRE) motif located within 2kb and 200bp upstream and downstream, respectively, from the TSS of each gig gene. Red, orange and grey colors inside the box indicate ISRE motifs associated with differentially regulated (dap) regions, ISRE motifs without association to dap regions, and lack of ISRE motifs, respectively. Created in BioRender. Manousi, D. (2025) https://BioRender.com/ubfasd1

In contrast, gig2 genes exhibited relatively more conserved expression patterns, particularly among WGD-derived paralogs (**Figure 6**). Members of the gig2A clade, most of which are located in LORe regions of ssa04, were generally upregulated in response to poly(I:C) stimulation and viral infection. Exceptions constituted the genes *gig2A_ssa08a and gig2A_ssa08c*, which showed low baseline expression and minimal induction. Similarly, gig2B paralogs (*gig2B_ssa12a* and *gig2B_ssa02d*) showed interferon-induced upregulation in head kidney cells.

Expression responses among gig2C paralogs were more variable (**Figure 6**). While *gig2C_ssa02a* and *gig2C_ssa02b* exhibited little expression under any condition, *gig2C_ssa12c* and *gig2C_ssa24* showed detectable responses to infection, indicating that regulatory divergence may be emerging within this clade. However, given the limited number of gig2C genes, further data would be required to draw definitive conclusions about functional divergence in this group. Taken together, these results suggest that gene sequence similarity does not translate to similarity at the level of transcriptional regulation. While some closely related paralogs maintain similar expression patterns, others have very distinct expression patterns, potentially reflecting functional specialization or even differential adaptation to immune stimuli. These findings underscore the complex interplay between duplication history and gene regulatory evolution in shaping the *gig*-immune genes functional repertoire in salmonids.

### Cis-regulatory motif analysis highlights context-dependent regulation of *gig* genes

To further elucidate the mechanisms underlying transcriptional regulation of *gig* genes in Atlantic salmon, we performed a scan for transcription factor (TF) binding motifs. Given the established classification of *gig* genes as IFN-stimulated genes (Zhang and Gui 2004; Sun *et al*. 2014), we searched for interferon-stimulated response elements (ISREs) – canonical sequences that bind IFN-regulated TFs during antiviral responses-in close proximity to *gig* genes. Using ISRE motifs with known impact in salmonids (Collet and Secombes 2001), and a physical proximity threshold of 2000 bp upstream and 200 bp downstream the TSS of each gene, we identified 25 ISRE motifs in 13 *gig1* genes and 21 ISRE motifs in 12 *gig2* genes (**Supplementary Table S5**).

We next identified genomic regions that showed chromatin accessibility changes in response to either poly(I:C)-induced interferon activation or SAV3 infection (Materials and Methods), a hallmark of transcription factor binding (Collet and Secombes 2001). We identified ISREs overlapping with chromatin accessibility peaks that were differentially regulated upon poly(I:C)-induced interferon activation. This allowed us to group *gig* genes into three categories: (i) genes obtaining nearby ISRE motifs associated with virally induced chromatin accessibility changes, (ii) genes obtaining nearby ISRE motifs without association to chromatin accessibility changes, and (iii) genes lacking nearby ISRE motifs (**Figure 6**). Comparison of expression profiles revealed a general trend: genes in class (i) tended to show higher expression under immune stimulation (**Figure 7**), but this difference was only significant for *gig1* genes (Mann–Whitney U test, *gig1*: p = 0.012, *gig2*: p = 0.37).

**Figure 7.**
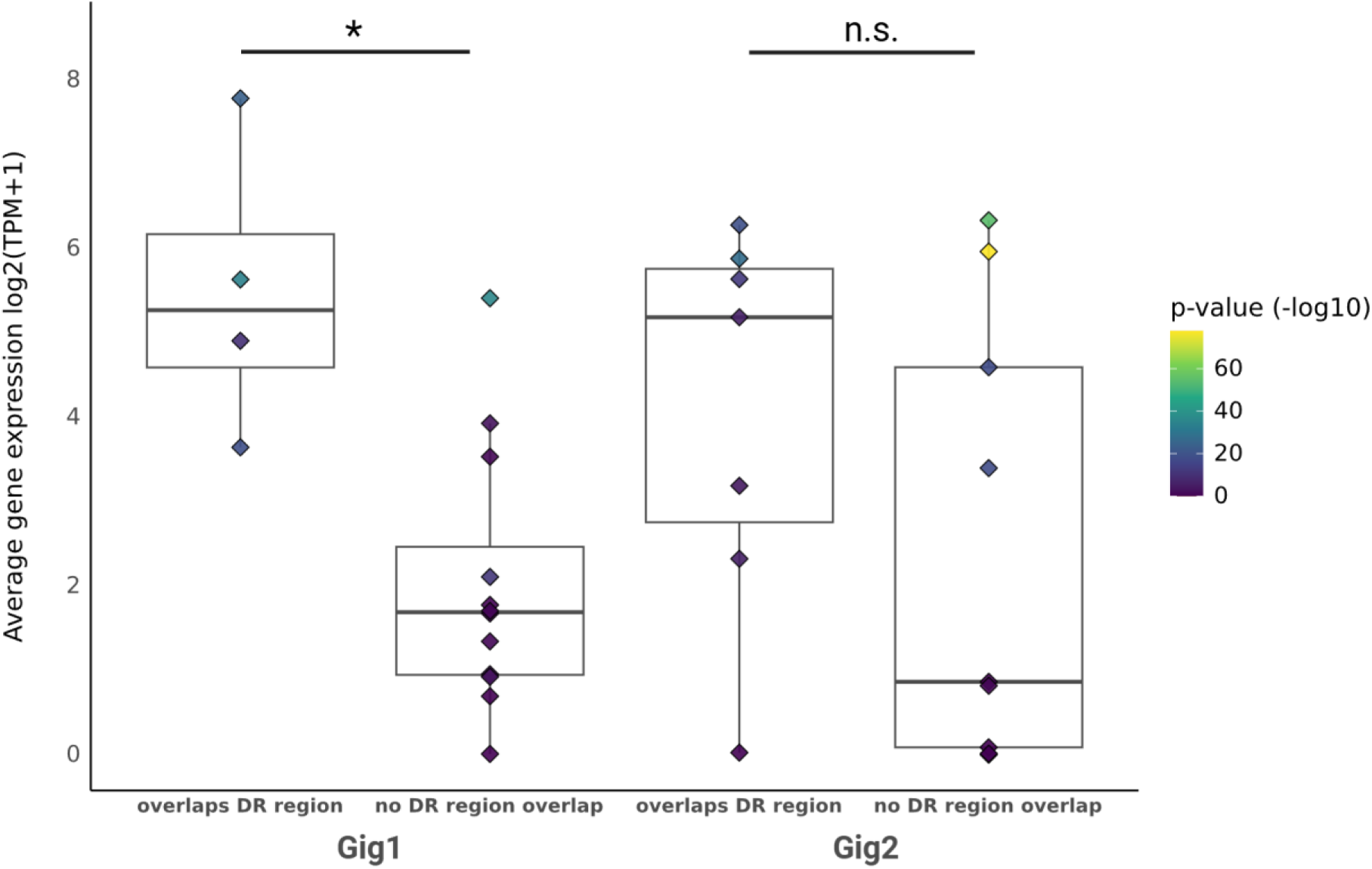
Distribution of genes with regulatory machinery near the TSS region. Different boxplots indicate genes with or without differentially regulated (DR) machinery during response to poly(I:C) viral mimic. Additionally, colored points represent differentially or non-differentially expressed gig1 and gig2 genes with their respective average expression shown in Y axis. Asterisks indicate the statistical significance (p-value < 0.05) in comparison of average gene expression and DR regulator for each of the two gig families. Created in BioRender. Manousi, D. (2025) https://BioRender.com/ikkn1ux

Interestingly, exceptions to this pattern were observed in tandemly duplicated genes. For example, *gig1B_ssa03a* harbored a transcriptionally active ISRE and showed upregulation, while its tandem duplicates, *gig1B_ssa03b* and *gig1B_ssa03c*, which were also upregulated lacked nearby ISREs (**Figure 7**). These findings suggest that cis-regulatory elements may exert effects across multiple adjacent paralogs or that alternative, uncharacterized motifs contribute to *gig* gene expression. Such context-specific transcriptional activity underscores the complexity of regulatory evolution in duplicated genomic regions.

Together, these results demonstrate that the transcriptional regulation of *gig* genes reflects both conserved IFN-driven control and divergent, context-dependent mechanisms.

## Discussion

### Evolutionary origin and diversification of gig genes

The *gig* gene families constitute a group of interferon-stimulated genes (ISGs) that are absent in mammals but conserved and expanded in fish and other non-amniote vertebrates (Zhang and Gui 2004; Zhang *et al*. 2013; An *et al*. 2025). These genes exhibit transcriptional activation during antiviral responses, indicating their important roles in vertebrate immune evolution (Sun *et al*. 2013; Gao *et al*. 2017; Madushani *et al*. 2021). In Atlantic salmon, a species of additional importance for the aquaculture industry, *gig* genes have shown involvement in antiviral responses (Krasnov *et al*. 2011, 2021) while recent studies suggested that tandemly duplicated *gig1* genes, in particular, may play key roles in antiviral response against important aquaculture diseases (Manousi *et al*. 2025). These findings suggest the necessity to further explore the evolutionary and functional divergence of *gig* genes, not only to deepen our understanding of immune evolution in teleosts but also to inform disease management strategies and sustainability efforts within aquaculture.

Our comparative analyses revealed that while *gig* genes are upregulated in response to viral challenges, the *gig1* and *gig2* families have evolved along distinct evolutionary trajectories. In his study, phylogenetic analysis across teleost species with additional whole-genome duplications, including Atlantic salmon and common carp, revealed a complex interplay between gene duplication and gene regulatory divergence in shaping gig immune gene repertoires. Interestingly, the divergent evolutionary trajectories of *gig1* genes between salmonids and cyprinids may reflect lineage-specific responses to ecological niches and pathogen pressures. While several salmonid species (including those used in our study) face the challenges of anadromous life cycles and varied pathogen exposures across freshwater and marine environments (Vollset *et al*. 2021), common carp inhabit more stable freshwater habitats (Banet *et al*. 2022), with distinct immune challenges. The observed differences in expansion, retention, and divergence of *gig* genes might therefore reflect selection pressures from lineage specific highlight the importance of lineage-specific host ecology and environment.

The phylogenetic and sequence conservation analyses in our study identified higher sequence divergence of *gig1* genes –reflected in the proportion of retained amino acids following sequence reliability filtering-as well as substantial length variation of the *gig1* protein sequence and predicted protein-coding domain between phylogenetic clades. These results suggest possibly broader functional divergence of *gig1* genes compared to the *gig2* family. Still, both gene families exhibited extensive sequence divergence and high gene gain and loss turnover rates, with frequent gene gains and losses shaping their evolutionary histories. The observed evolutionary patterns align with previous findings in *gig* genes (Zhang *et al*. 2013; Sun *et al*. 2014; An *et al*. 2025) but also with patterns of rapid gene birth-and-death evolution observed in other IFN-related genes (Redmond *et al*. 2019). The observed sequence diversity and difficulty in reconstruction of clear phylogenetic relationships within both *gig* families in the present study is in line with the idea that these genes are evolving fast under strong selective pressures driven by host-pathogen interactions (Shultz and Sackton 2019).

### Structural evolution of *gig1* and *gig2* genes in Atlantic Salmon

In our focused analysis of *gig* evolution in Atlantic salmon, functional domain analysis identified family-specific domains, consistent with previous studies in other teleosts (Zhang and Gui 2004; Sun *et al*. 2013). Interestingly, while earlier work suggested that *gig1* proteins primarily reside in the cytoplasm (Sun *et al*. 2014), our analysis revealed that a subset of Atlantic salmon *gig1* genes carry membrane-associated domains. Although only few of these membrane-associated *gig1* paralogs exhibited significant transcriptional responses to viral infection (**Figure 6**), acquisition of the particular predicted domain across gig paralogs is hinting at potential divergence in subcellular localization.

Additionally, repeat elements were frequently found across *gig1* and *gig2* paralogs. Large repeats in certain *gig1* genes were associated with apparent gaps in functional domain prediction, although manual annotation confirmed the integrity of the coding sequences. In *gig2*, repeat expansions coincided with duplicated functional domains, particularly in *gig2_ssa08b* and *gig2_ssa12b*, while shorter tandem repeats contributed to glutamate (E)-rich disorder regions in *gig2_ssa02d* and *gig2_ssa12a* (**Figure 5**). Repeat-driven genomic rearrangements are known to influence protein stability and plasticity, facilitating functional diversification (Rivera and Swanson 2022). For immunity in particular, the presence of repeat content in the amino acid sequence can increase immunogenetic plasticity, promoting functional diversification over short evolutionary timescales, which is an important factor in the arms race between hosts and pathogens (Teekas *et al*. 2022) In our dataset, repeat-containing *gig* genes often exhibited low expression and tissue-specific responses, suggesting that such signatures of structural domain divergence may be related to adaptive specialization –to additional roles or be a signature of pseudogenization processes following gene family expansion.

### Transcriptional Regulation and Functional Divergence of *gig* genes in Atlantic salmon

We identified presence of IFN-stimulated response elements (ISREs) in 25 out of 32 *gig* genes, supporting their predicted role in IFN-dependent immune regulation. However, only a subset of *gig1* and *gig2* paralogs exhibited regulatory activity during viral infection, concordant with the idea that transcriptional control, rather than simple promoter presence, modulates functional relevance (Ray-Jones and Spivakov 2021). Consistent with their classification as interferon-stimulated genes, *gig* genes showed strong responses to both poly(I:C)-induced (interferon-mediated) immune activation and SAV3 infection in heart and head kidney cells and tissues. While most paralogs exhibited similar expression across different infection conditions, a small subset showed distinct regulatory responses, suggesting functional divergence among retained copies. Interestingly, tandemly duplicated *gig* genes did not consistently share expression patterns, implying that regulatory elements may be exchanged between physically adjacent paralogs. This pattern aligns with previous findings in antiviral genes, where tandem duplication has led to transcriptional interference or regulatory divergence over evolutionary time (Krasnov *et al*. 2021). At the same time, correspondence of *gig* expression between IFN-stimulated cells and tissue samples indicates the possibility of studying the functional divergence of *gig* genes in *in vitro* models, thus reducing the need for conduction of infectious challenges and consequent sacrifice of live fish.

Despite their established role as IFN-responsive genes, earlier studies in grass carp suggested that *gig* genes might also be activated through an IFN-independent pathway (Sun *et al*. 2013). Our findings in Atlantic salmon suggest that while *gig* genes largely function within an IFN-dependent framework, regulatory divergence may have led to paralog-specific tuning of immune responsiveness. Notably, some *gig* paralogs exhibited consistently low expression, raising the possibility of functional redundancy or pseudogenization following multiple rounds of whole-genome and segmental duplication, supporting the idea of a combination of selection and drift shaping the evolution of the *gig* gene family. Such patterns align with recent studies on antifreeze proteins (AFPIs) in fish, which were shown to have originated from non-coding sequences of a seemingly unrelated *gig2* gene (Graham *et al*. 2022; Rives *et al*. 2024). This provides a marked example of how functionally redundant paralogs can be retained as raw material for evolutionary innovation, reinforcing the idea that immune genes frequently undergo repurposing under selective pressures. Still, further analysis is required, potentially through application of targeted gene editing approaches in order to identify the regulatory mechanisms and functional impact of *gig* paralogs.

### Decoupling of ISRE-motif presence and transcriptional regulation

Our findings in Atlantic salmon suggest that *gig* genes in Atlantic salmon largely function within an IFN-dependent framework. Nineteen out of 32 gig-genes displayed some form of experimental transcriptional immune activation, and sixteen of these (84%) also had ISRE-elements annotated in their promoters. Nevertheless, 8 gene genes had ISRE-motifs but showed no transcriptional response while 3 genes had no ISRE-motif but showed transcriptional response (**Figure 6**). Finally, five of the immune-responding genes did have ISRE-motifs but no evidence for chromatin accessibility changes, i.e. TF binding, overlapping the motif. Some of these discrepancies are likely to be associated with technical aspects of the functional genomics experiments leading to false negatives; for example low power to detect small effect sizes or species specific alternative variants of ISRE-motifs. However, these discrepancies could also be due to real biological phenomena.

Several mechanisms could explain the lack of correlation between ISRE-motif presence and transcriptional regulation. Despite their established role as IFN-responsive genes, earlier studies in grass carp suggested that *gig* genes might also be activated through an IFN-independent pathway (Sun *et al*. 2013). This could also be the case for the three gig-genes in Atlantic salmon that responded to immune challenges across several independent experiments but did not have ISRE-motifs present in their promoter sequences. Another possibility is that these three ISRE-lacking genes were in fact transcriptionally responsive through ISRE-motif binding but that the motifs were residing in distant enhancers (Leviyang 2021) and hence not identified in our bioinformatics approach. Finally, several tandemly duplicated *gig* genes in our study have the same annotation of ISRE-promoter element but did not consistently share expression patterns. This pattern could be explained by transcriptional interference (Proudfoot 1986; Shearwin *et al*. 2005), a process which has previously been proposed to lead to transcriptional interference between the tandemly duplicated genes (Loker and Mann 2022).

## Conclusion

In conclusion, our study reveals that *gig* genes have undergone extensive and lineage-specific diversification in teleosts, shaped by whole-genome duplications, tandem expansions, and repeat-driven structural changes. In Atlantic salmon, we show that *gig1* and *gig2* families follow distinct evolutionary trajectories, with marked differences in sequence conservation, domain structure, and transcriptional regulation. Notably, our data indicate that regulatory divergence, including decoupling between ISRE motifs and transcriptional responses, is a key feature of gig gene evolution. Such patterns reflect a complex interplay between selective pressures and genomic context, contributing to functional diversification among paralogs.

These findings deepen our understanding of immune gene evolution in aquatic vertebrates and reveal the potential of gig genes as targets for functional studies and applied breeding strategies. By leveraging the evolutionary plasticity of these IFN-stimulated genes, aquaculture programs may develop more resilient populations with enhanced resistance to viral pathogens, contributing to the long-term sustainability of aquaculture.

## Supplementary Figures

**Supplementary Figure 1.**
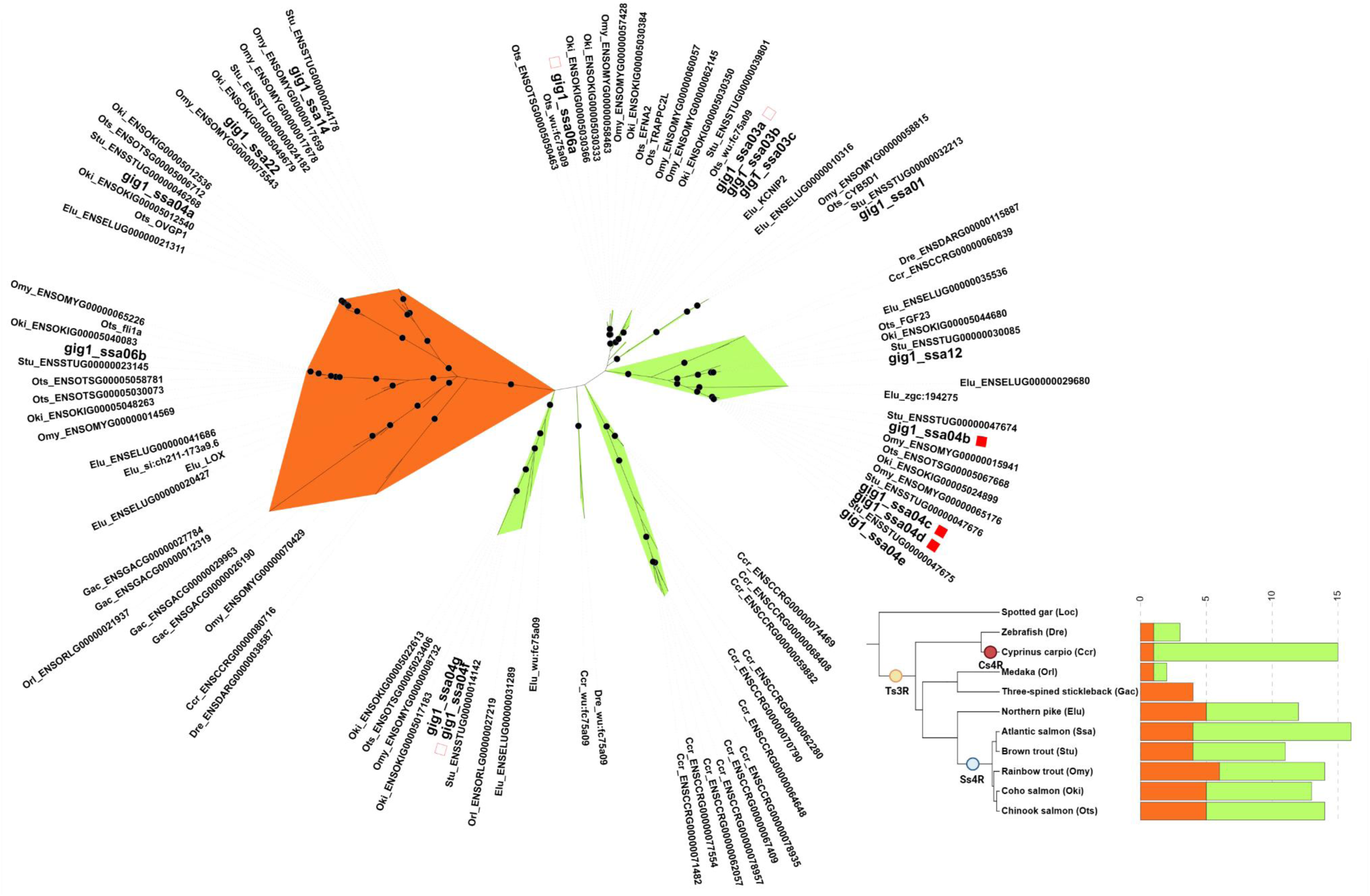
Unrooted phylogenetic tree of gig1 genes detected across 11 species. Ortholog genes across species were retrieved from the Ensembl annotation gene gain/loss trees. Gene names were retrieved using Biomart (Supplementary Table S1). The protein translation of the coding sequence for the longest transcript of filtered genes was aligned using GUIDANCE2 and unreliable residues were removed based on the default threshold of 0.93. A phylogenetic tree was built in IQ-TREE v2.3.3 using the filtered alignment and 1000 replicates to calculate bootstrap values. The ‘Model Finder’ option was additionally applied to identify and use the best-fit substitution model for the tree. Sequence divergence within the gig1 family suggests one possible split in the middle of the tree forming two separate clades, hereby named gig1A (orange) and gig1B (green). Red squares indicate repeat content in the coding sequence of the respective genes with filled squares indicating large repeats and empty squares showing small repeats. The number of gig1 genes across species are shown in the bottom right bar chart where colored ranges indicate the respective gene clusters and yellow, red and blue annotations represent the teleost-specific (Ts3R), carp-specific (Cs4R) and salmonid-specific (Ss4R) whole genome duplication events, respectively. Created in BioRender. Manousi, D. (2025) https://BioRender.com/y2vfiz6

**Supplementary Figure 2.**
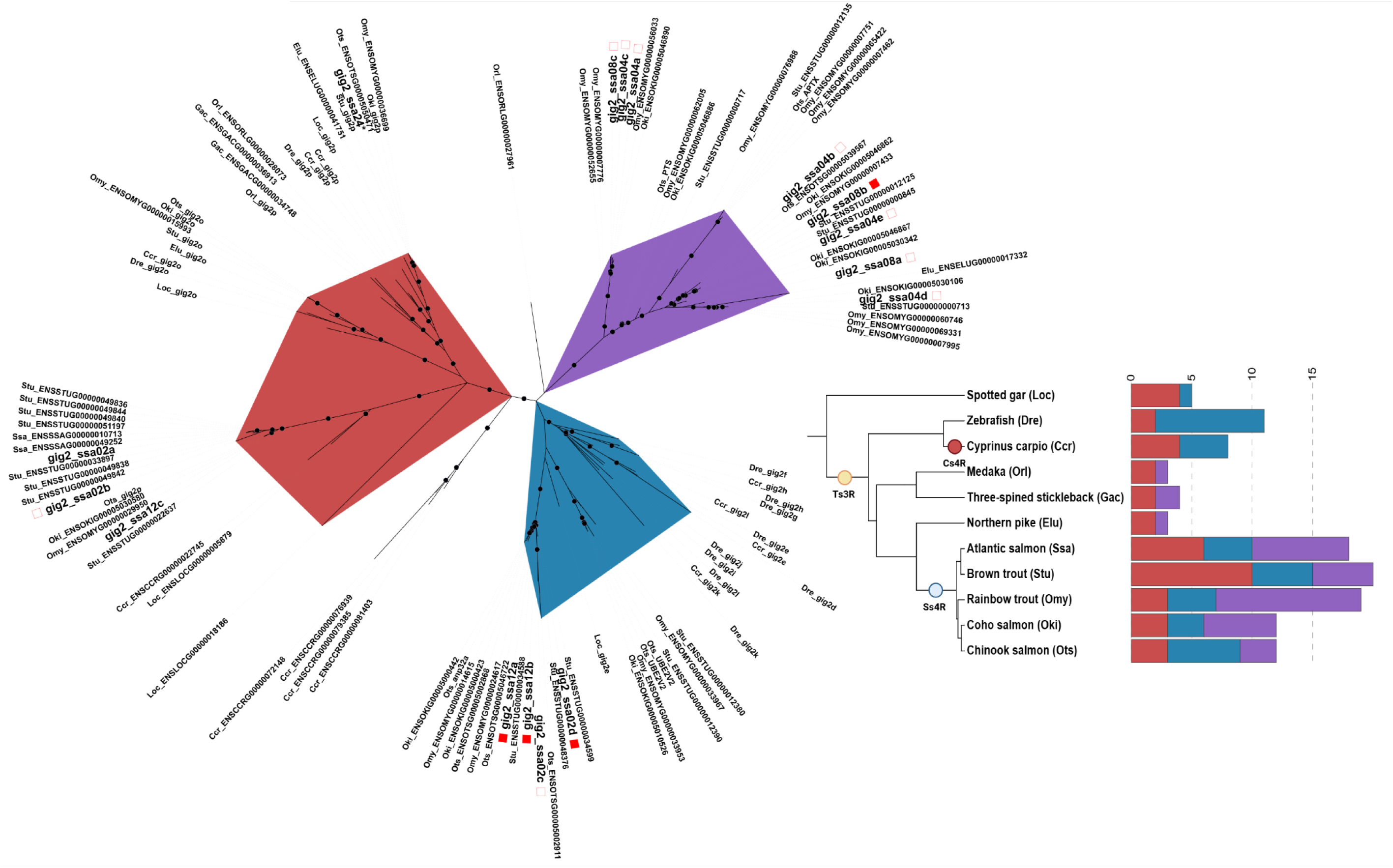
Unrooted phylogenetic tree of gig2 genes detected across 11 species. Ortholog genes across species were retrieved from the Ensembl annotation gene gain/loss trees whereas gene names were retrieved using Biomart. The protein translation of the coding sequence for the longest transcript of each gene was aligned using GUIDANCE2 and unreliable residues were removed based on the default reliability threshold of 0.93. A phylogenetic tree was built in IQ-TREE v2.3.3 using the filtered alignments and 1000 replicates to calculate bootstrap values. In addition, the ‘Model Finder’ option was applied to identify the best-fit convergence model for the tree. Sequence divergence within the gig2 family suggested three possible splits in the middle of the tree forming three separate clades, hereby named gig2A (red) and gig2B (blue) and gig2C (purple). Red squares indicate repeat content in the coding sequence with filled squares indicating large repeats and empty squares showing small repeats. The number of gig2 genes across species are shown in the bar chart where colored ranges indicate the respective gene clusters and yellow, red and blue tree annotations represent the teleost-specific (Ts3R), carp-specific (Cs4R) and salmonid-specific (Ss4R) whole genome duplication events, respectively. Created in BioRender. Manousi, D. (2025) https://BioRender.com/8narlgo

**Supplementary Figure 3.**
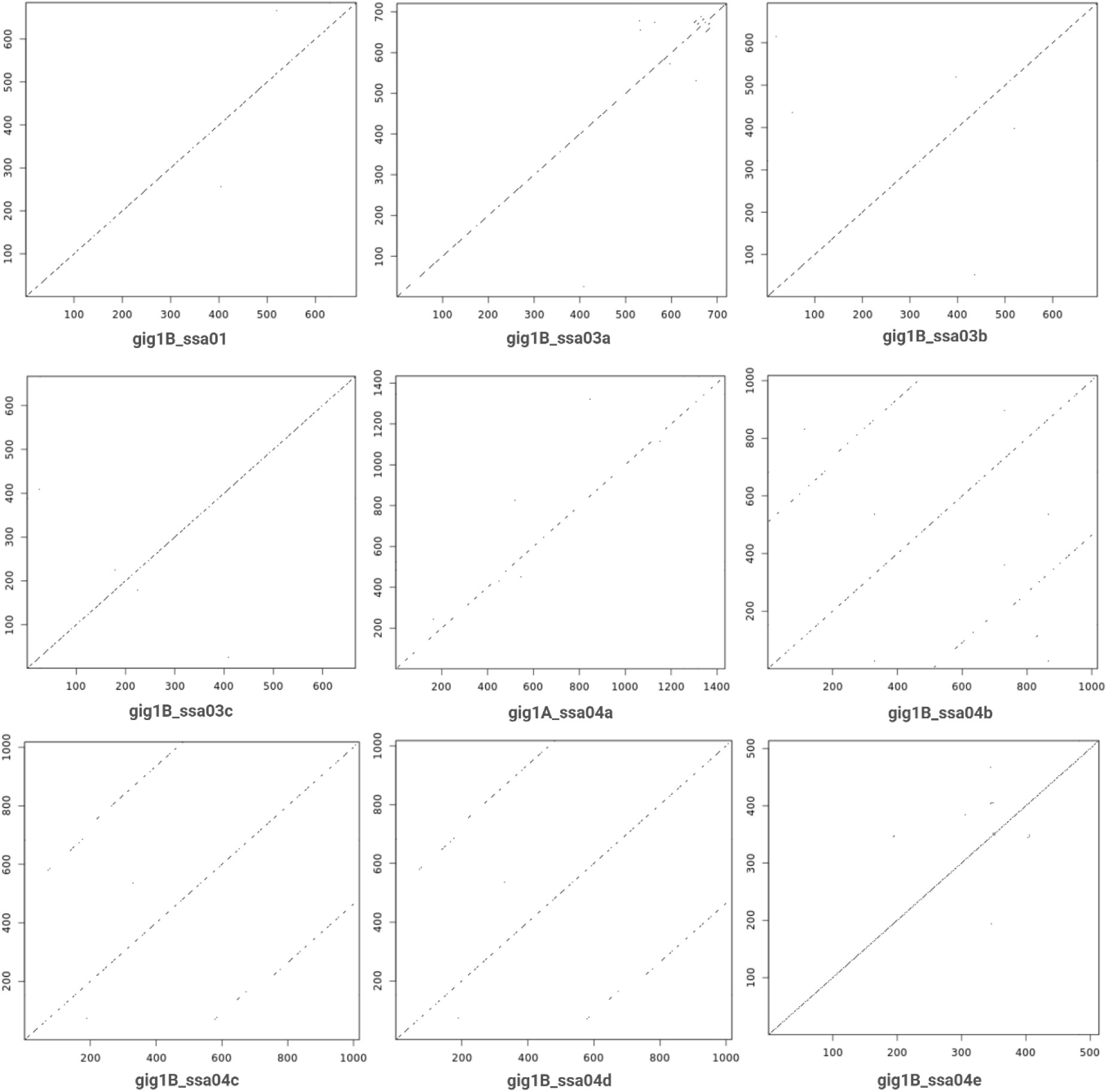
Pairwise comparison of the coding sequence of gig1 genes. To identify repeat content within gig paralogs, the nucleotide sequences were analyzed using a self-comparison approach to detect internal sequence repetitions. Created in BioRender. Manousi, D. (2025) https://BioRender.com/ixs4lqw.

**Supplementary Figure 4.**
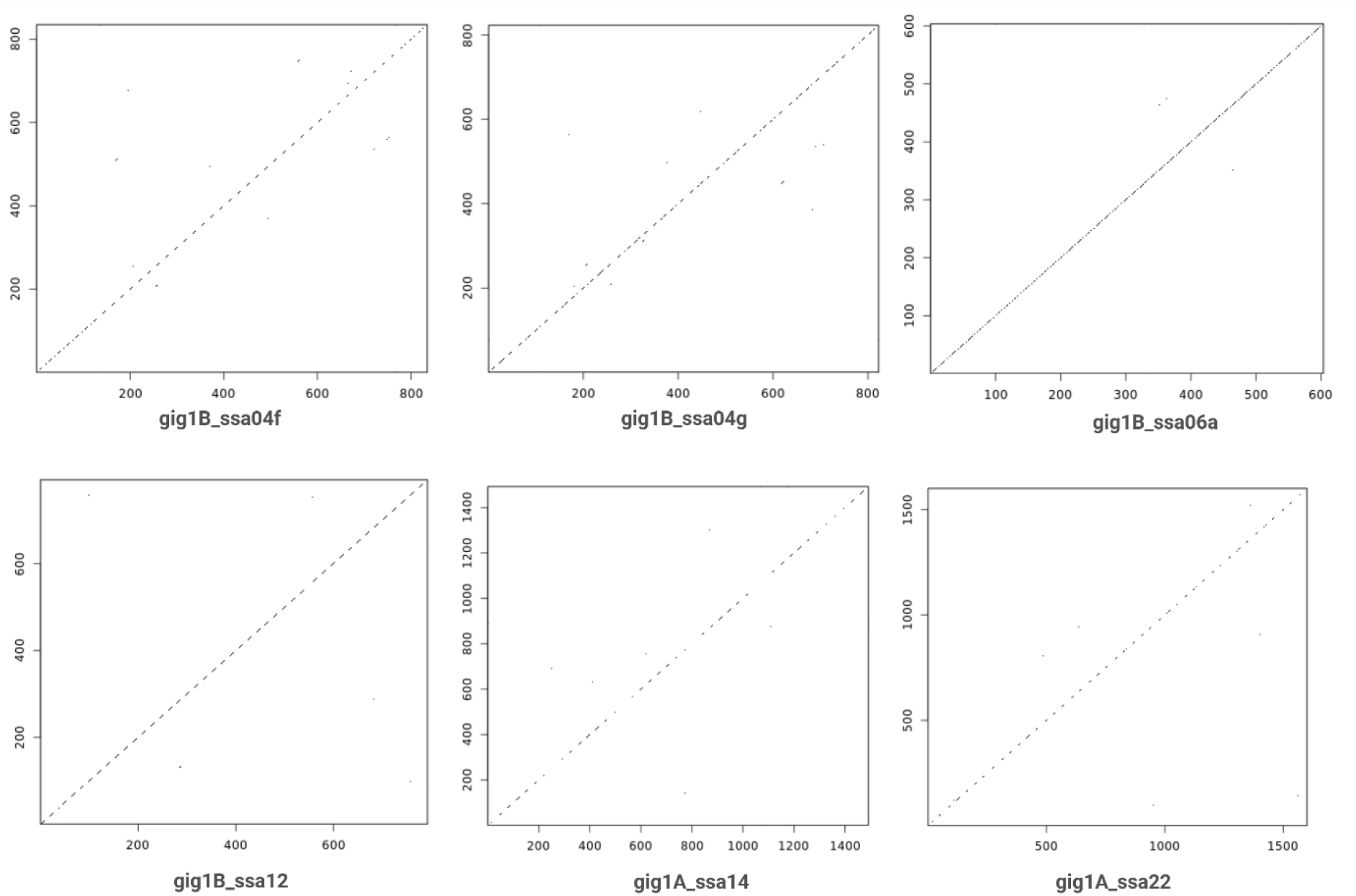
Pairwise comparison of the coding sequence of gig1 genes. To identify repeat content within gig paralogs, the nucleotide sequences were analyzed using a self-comparison approach to detect internal sequence repetitions. Created in BioRender. Manousi, D. (2025) https://BioRender.com/zscon0j.

**Supplementary Figure 5.**
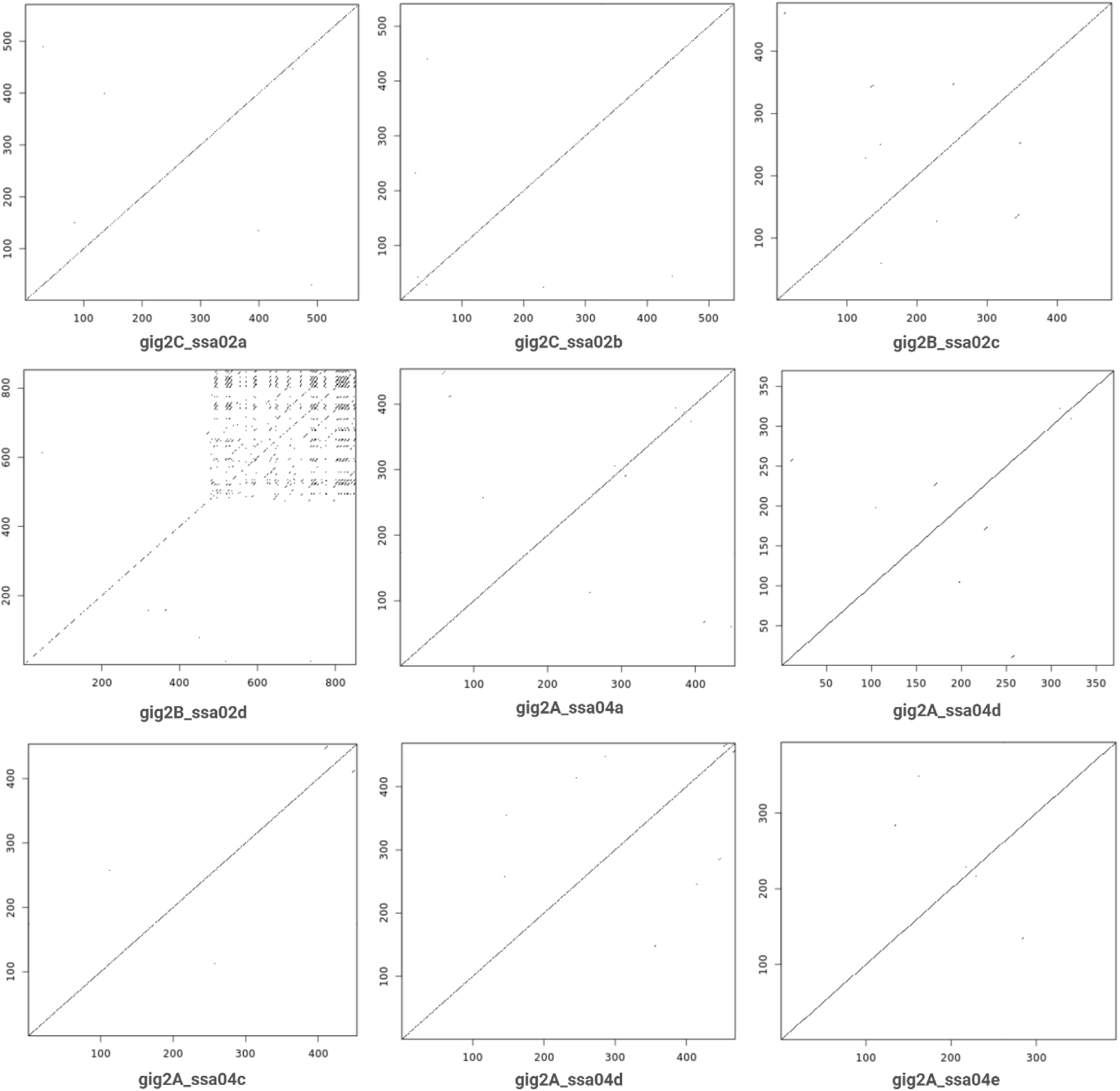
Pairwise comparison of the coding sequence of gig2 genes. To identify repeat content within gig paralogs, the nucleotide sequences were analyzed using a self-comparison approach to detect internal sequence repetitions. Created in BioRender. Manousi, D. (2025) https://BioRender.com/r977e2m

**Supplementary Figure 6.**
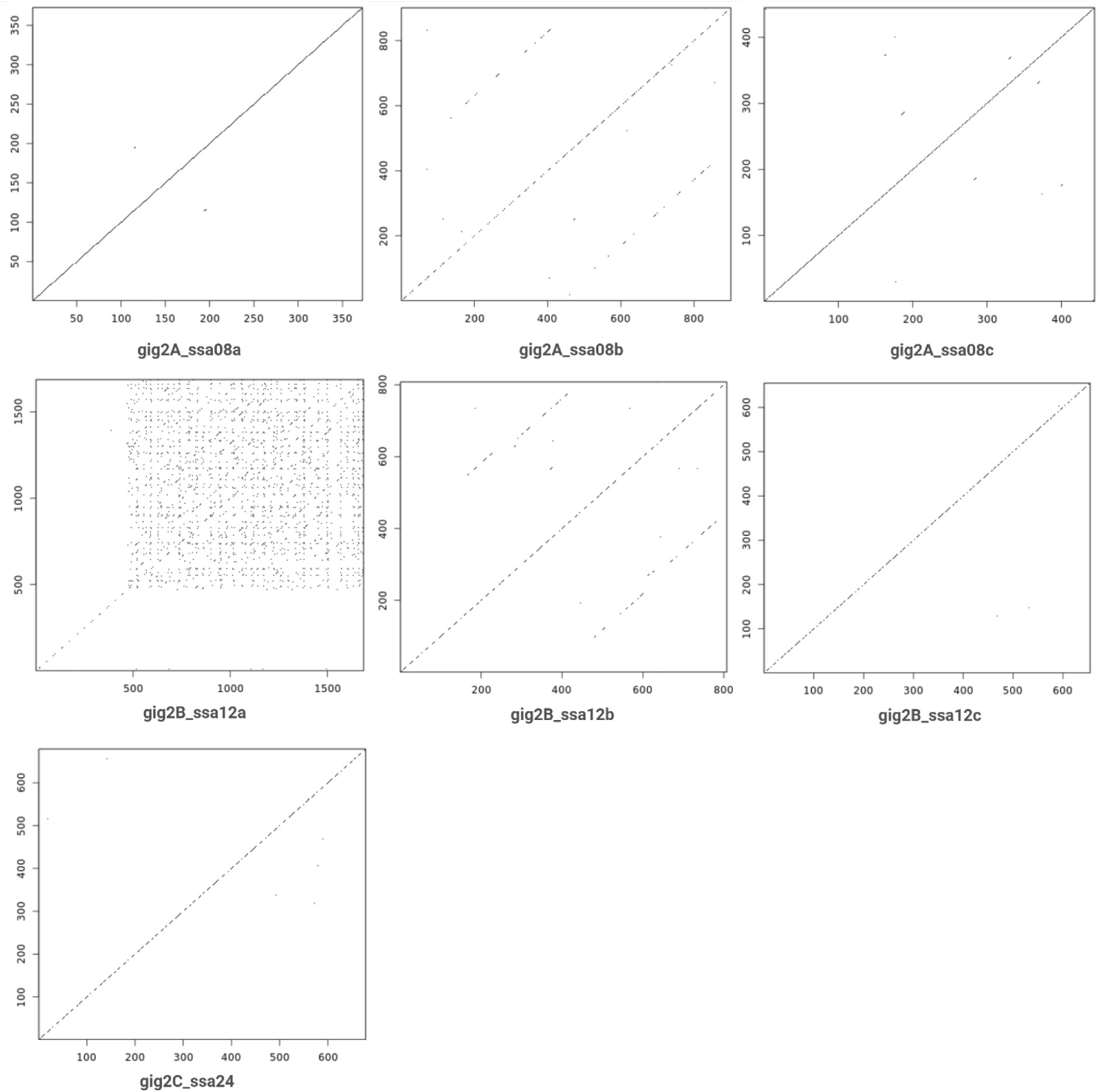
Pairwise comparison of the coding sequence of gig2 genes. To identify repeat content within gig paralogs, the nucleotide sequences were analyzed using a self-comparison approach to detect internal sequence repetitions. Created in BioRender. Manousi, D. (2025) https://BioRender.com/79g6wsd

## Supplementary Tables

### File S1: Manousi_gig_supplementary_tables.xlsx

*Table S1. Gig genes retrieved from 11 selected aquatic species were retrieved from the Ensembl database (version 112). Columns in the table represent the Ensembl gene ID for each species as well as the gene name and description provided by Ensembl. In addition, the gene ID and description provided from the NCBI genome database is provided where possible. Gene names, IDs and descriptions were obtained using the Biomart service in Ensembl (version 112)*.

*Table S2. Predicted functional features using the InterPro database and the amino acid translation of gig1 and gig2 genes across 11 species. The top section displays functional domains predicted across all species, whereas the bottom section focuses on domains identified in Atlantic salmon gig paralogs. Tables contain the name of predicted domains, the database from each they were retrieved, as well as the gig family and the abundancy each domain was discovered. Finally, Functional clusters denote the classification used in this analysis to cluster together similar functional domains retrieved from multiple databases*.

*Table S3. Comparison of the physical overlap of Atlantic salmon gig genes and chromosomal regions of either early or recent rediploidization timing. Synteny block information were retrieved from the Salmobase database (*https://salmobase.org/*), indicating duplicate chromosomal regions of early (Ancestral Ohnolog Resolution, AORe) or recent (Lineage-specific Ohnolog Resolution, LORe) rediploidization. The physical positions of gig genes were then compared against syntenic block to identify gig genes overlapping AORe/LORe regions*.

*Table S4. Gig gene detected in Atlantic salmon. The table shows all gig1 and gig2 genes identified in Atlantic salmon using the Ensembl database (version 112). Nomenclature of each gene was given based on their respective phylogenetic clade and physical position in the salmonid genome, shown in columns 3-6. To verify the structure of each gene, short and long transcriptomics data were visualized together with the Atlantic salmon functional annotation on IGV, manually annotating each gene. Where impossible to visually annotate, provided in column 8, the structure of the gene was based on the publicly available functional annotation*.

*Table S5. Table of detected Interferon stimulation response elements (ISRE) motifs in close proximity to gig genes. Briefly, motif detection was performed using ISRE sequences with known antiviral impact in salmonids (Collet and Secombes 2001) and a physical proximity threshold of 2000 bp upstream and 200 bp downstream the TSS region of each gene. Columns on the table show the respective genes and detected motif sequences within the defined physical threshold from the TSS. In addition, motifs are characterized as differentially regulated based on overlap of the sequence with differentially regulated ATAC-seq and CHIP-seq peaks after infection of head kidney samples with the poly(I:C) viral mimic*.

## Data Availability

Raw short-read transcriptomics data related to the SAV3 infectious cohabitation challenge is available on the European Nucleotide archive under accession number PRJEB85594. Raw short-read transcriptomics data related to the SAV3 survival challenge is available on the European Nucleotide archive under accession number PRJEB85594. Raw material for the analysis of the Atlantic salmon regulatory landscape is available on the European Nucleotide Archive under accession numbers PRJEB50077 and PRJEB56698 for ATAC-seq and ChIP-seq information, respectively. Raw short-read transcriptomics data related to the poly(I:C) challenge of Atlantic salmon head kidney cells and tissue are available on the European Nucleotide archive under accession number PRJEB50076. All scripts are available at https://github.com/DomManou/Gig_gene_divergence.

## Acknowledgements

We acknowledge the use of the Orion computing cluster at the Norwegian University of Life Sciences (NMBU) as well as the use of BioRender for the editing and visualization of all figures and supplementary figures in the manuscript.

